# Membrane Proteome Remodeling in Female APP Mice Following Muscarinic Acetylcholine Receptor M1 Modulation Revealed by Peptidisc Enabled DIA-MS

**DOI:** 10.64898/2026.02.24.707828

**Authors:** Ashim Bhattacharya, Frank Antony, Hiroyuki Aoki, Mohan Babu, Stephen S. G. Ferguson, Khaled S. Abd-Elrahman, Franck Duong van Hoa

## Abstract

Alzheimer’s disease (AD) is linked to profound dysregulation of membrane-embedded and membrane-associated proteins that govern amyloid processing, synaptic signaling, and neuronal communication. Yet most proteomic analyses prioritize soluble fractions, resulting in systematic underrepresentation of integral membrane proteins and limited access to disease-relevant membrane pathways. Here, we use a membrane-mimetic, data-independent acquisition proteomic workflow to define disease- and drug-induced remodeling of the cortical membrane proteome in an APP mouse model of Alzheimer’s disease. Female B6C3F1/J mice were aged to 9 months and treated for 8 weeks with or without the M1 muscarinic acetylcholine receptor positive allosteric modulator VU0486846. APP pathology drove a pronounced, genotype-specific remodeling of the membrane proteome, with enrichment of multiple membrane proteins linked to AD, including RyR2, PLD3, ITM2C, and CNTNAP2. Wild-type mice cortical membranes were instead enriched for membrane proteins involved in axon guidance and synaptic organization, such as EPHA5 and ROBO2. In contrast, activation of M1 using the VU0486846 produced minimal membrane proteome changes in wild-type mice but selectively enriched proteins involved in neuronal trafficking and synaptic plasticity in APP mice, including SORCS2, PLXND1, and CADM1. Together, these findings demonstrate that AD-associated proteomic remodeling is strongly concentrated at the membrane level and that M1 receptor activation preferentially engages disease-altered membrane networks rather than inducing widespread proteomic changes. This work establishes peptidisc-enabled membrane proteomics as a powerful approach for identifying membrane-associated biomarkers and evaluating therapeutic target engagement in AD.

Graphical abstract

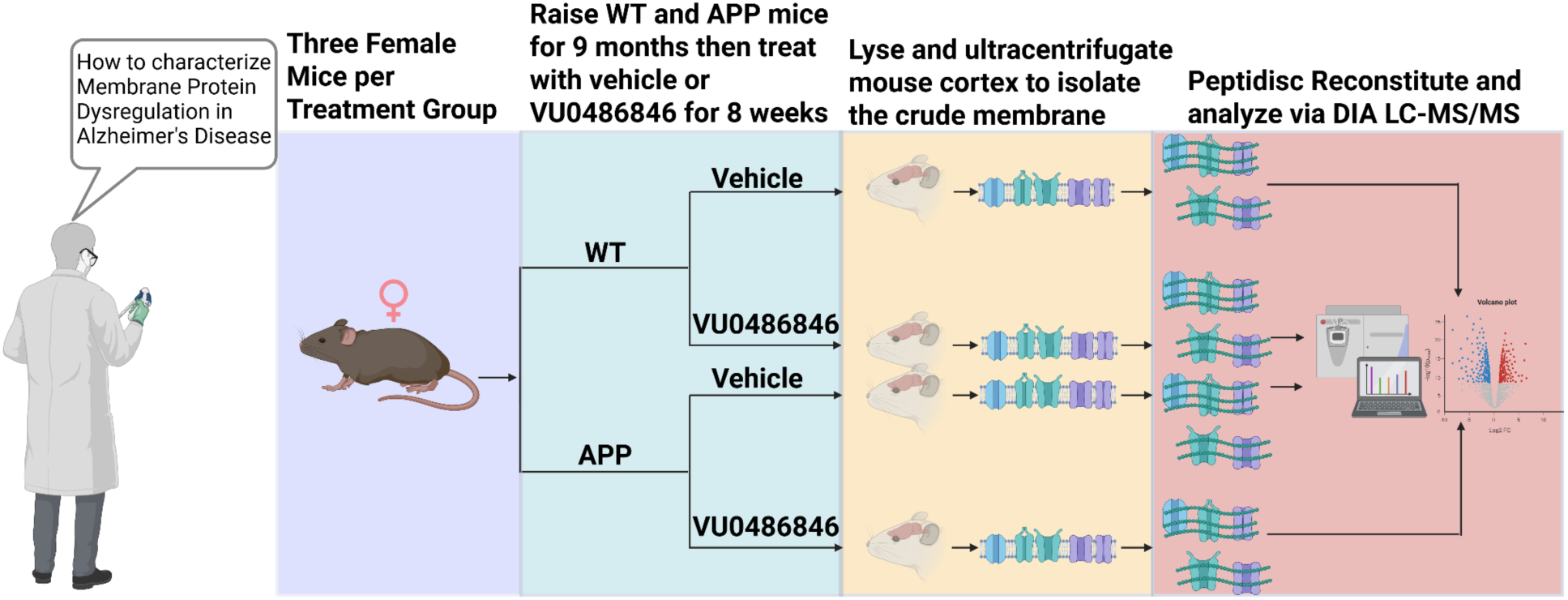

## Introduction

Alzheimer’s disease (AD) is the most common neurodegenerative disorder and the leading cause of dementia worldwide, ranking as the seventh leading cause of death. Despite decades of research, there is no curative therapy (1). Accordingly, AD research constitutes a major focus of contemporary biomedical science, with the U.S. National Institutes of Health dedicating approximately $3.9 billion per year (2), or ∼8% of its total budget (3), to Alzheimer’s and related dementia research. To this end, there is a clear need for novel biological insights and methodological advances to deepen our understanding of AD pathology and to enable the discovery of robust biomarkers and therapeutic targets.

Transcriptomic approaches have been applied extensively to AD, providing valuable insight into disease-associated gene expression programs (4), whereas proteomic analyses have historically been less frequently performed, in part due to technical complexity and sample requirements (5,6). As it has become increasingly clear that RNA abundance does not reliably predict protein abundance, modification state, or localization, there has been growing interest in proteomic approaches to more directly interrogate disease-relevant molecular changes (7). To date, proteomic studies of AD have relied on clarified tissue lysates generated by centrifugation at a moderate/low speed step that removes cellular debris and membrane fractions and enriches the soluble proteome (6,8–10). Consequently, soluble-enriched proteomic workflows are inherently biased toward non-membrane proteins, leaving membrane-embedded receptors, transporters, and signaling proteins, central to synaptic function, therapeutic targeting, and disease pathology, poorly resolved (11).

To address the long-standing bias of mass spectrometry (MS)–based proteomics toward soluble proteins, we recently performed a systematic comparative analysis to identify the optimal strategy for membrane proteome enrichment and detection of integral membrane protein (IMP)–level dysregulation. Specifically, we benchmarked three membrane mimetic–based workflows (Peptidisc, nanodisc, and SMA2000) alongside four detergent clean-up–based methods (SP3, SP4, FASP, and S-Trap) in the context of healthy vs obese mouse liver membrane proteomics. Through this analysis, the Peptidisc membrane mimetic emerged as the most robust approach, exhibiting the lowest variability across replicates and the highest sensitivity for detecting IMP-level dysregulation in the context of a mouse liver metabolic disease model (12).

In this study, we applied a peptidisc-based membrane proteome profiling strategy to a well-validated APP transgenic mouse model of AD. These mice express familial AD–associated APP mutations in combination with presenilin-1 or presenilin-2 (PSEN1/2), driving amyloid-β overproduction, plaque deposition, and cognitive impairment that recapitulate key features of human AD pathology (13). Given the central role of muscarinic acetylcholine receptor M1 (M1 mAChR) signaling in synaptic function and cognition, and its established relevance to AD, we further examined how pharmacological modulation of M1 signaling reshapes the disease-associated membrane proteome.

To selectively enhance M1 mAChR signaling while avoiding the cholinergic side effects and receptor desensitization associated with direct agonists (14), we employed the M1-selective positive allosteric modulator VU0486846 (15,16). Previous studies have demonstrated that VU0486846 improves cognitive performance in APP mice, as assessed by novel object recognition and Morris water maze tasks, in both male and female animals (16,17). However, VU0486846 reduced amyloid-β plaque burden only in female mice, with no comparable effect observed in males, indicating a pronounced sex-dependent response (15,16). This finding is consistent with prior reports of sex-specific differences in M1 mAChR expression and signaling in the brain (18). In light of this dichotomy, we focused our analysis on female APP mice to maximize the sensitivity and reproducibility of detecting M1-dependent membrane proteome remodeling associated with AD pathology.

Using peptidisc-enabled DIA mass spectrometry, we detected robust APP overexpression and dysregulation of multiple established AD-associated integral membrane proteins in APP mice. In contrast, treatment with VU0486846 produced minimal membrane proteome changes in wild-type (WT) mice but induced substantial remodeling in APP mice, including increased abundance of proteins involved in neuronal trafficking and synaptic plasticity. Together, these results show that peptidisc-based membrane proteomics enables sensitive detection of membrane proteome dysregulation in AD mice as well as more subtle ligand-induced membrane-associated processes through which M1 mAChR activation mitigates AD pathology.

## Materials and Methods

### Materials

VU0486846((R)-4-(4-(1H-pyrazol-1-yl)benzyl)-N-((1S,2S)-2-hydroxycyclohexyl)-3,4-dihydro-2H-benzo[b][1,4]oxazine-2-carboxamide; compound ID 3271) was obtained from Axon Medchem. n-Dodecyl-β-D-maltoside (DDM) was purchased from Anatrace. His₆-tagged peptidiscs (purity >90%) were obtained from Peptidisc Biotech. Nickel nitrilotriacetic acid (Ni-NTA) chelating Sepharose resin was purchased from Qiagen. Complete protease inhibitor cocktail was obtained from Sigma. Amicon Ultra centrifugal filters (100 kDa molecular weight cut-off) were purchased from Millipore. MS-grade trypsin and DNase I were obtained from Thermo Fisher Scientific. Octadecyl C18 Empore disks were purchased from 3M, and Polygoprep 300-20 C18 powder was obtained from Macherey-Nagel. All other general reagents, including NaCl, Tris base, dithiothreitol (DTT), phenylmethylsulfonyl fluoride (PMSF), iodoacetamide (IAA), urea, organic solvents, and acids, were obtained from BioShop or Fisher Scientific Canada.

### Treatment and harvest of animal organs

The APP mouse model, VU0486846 treatment and *in vivo* characterization, including pathology, cytological analyses, neurogliosis, and behavioral outcomes, have been reported previously by our group (15,16,17). Briefly, mice were obtained from The Jackson Laboratory and bred to generate littermate-controlled female wild-type (B6C3F1/J, stock no. 100010) and APPswe/PSEN1ΔE9 (B6C3-Tg(APPswe/PSEN1ΔE9)85Dbo/J, stock no. 034829) cohorts.

Animals were group-housed (≥3 per cage), provided food and water *ad libitum*, and maintained on a 12 h light/12 h dark cycle at 24 °C. Female wildtype and APP/PS1 mice were aged to 9 months and were randomized and blindly-treated for 8 weeks with either vehicle (10% DMSO) or VU0486846 (10 mg/kg/day dissolved in 10% DMSO) in drinking water containing 1% sucrose and 0.5% tween. Water intake and drug concentration were calculated based on the average water consumption of 4 ml/day for a 25 g mouse (16). At the end of the 8-week treatment, mice were sacrificed, and their brains were collected. All animal experimental protocols were approved by the University of Ottawa Institutional Animal Care Committee and were in accordance with the Canadian Council of Animal Care guidelines.

### Tissue processing and preparation of the membrane fraction

Frozen tissue samples were thawed on ice, minced, and subsequently homogenized in hypotonic lysis buffer (10 mM Tris-HCl pH 7.4; 30 mM NaCl, and 1 mM EDTA, 1x cocktail protease inhibitor and 1 mM PMSF). The homogenate was further incubated for 10 min on ice in the presence of 10 mM MgCl2 and 50 μg/ml DNAse. The enlarged cell suspension was lysed using a French press with 3 passages at 500 pounds per square inch (psi). The cell lysate was centrifuged at 1,200 x g for 10 min at 4◦C to remove unbroken cells and nucleus fraction. The supernatant was collected and centrifuged at 5,000 x g for 10 min at 4◦C to remove the mitochondrial fraction, and the supernatant was ultracentrifuged at 110,000 x g for 45 min at 4◦C in a Beckman TLA110 rotor. The pellet was resuspended in 200 μL of TSG buffer (50 mM Tris, pH 7.9, 100 mM NaCl, 10% glycerol) to obtain a membrane fraction called “crude membrane” and stored at −80°C until use.

### Sample preparation for Peptidisc-based analysis

Crude membranes (∼1 mg) were resuspended in ice-cold TS buffer supplemented with 1% (w/v) DDM for 30 min at 4◦C with gentle shaking. The detergent extract was clarified by ultracentrifugation (110,000 x g for 15 min at 4◦C), and 1 mg aliquots (500 μl) were incubated with a 3-fold excess of His6-tagged Peptidiscs for 15 min at 4◦C. The sample was rapidly diluted to 5 mL in the TS buffer placed in a 100 kDa cutoff centrifugal filter. The sample was concentrated at 3,000 x g, 10 minutes to approximately 200 μl and diluted again to 5 ml, followed by another concentration step to approximately 200 μl. The reconstituted start Peptidisc library (4 mg total in 1 mL) was incubated with 60 μL of Ni-NTA chelating Sepharose for 1 hour at 4◦C with shaking. Following five thorough washes with TS buffer 1 mL each, the Peptidisc library was eluted in 150 μL of TS buffer containing 600 mM imidazole, recovering 1 mg at a concentration of ∼6.5 μg/μL. All fractions from the reconstitution through the purification steps were subjected to analysis by 15% SDS PAGE, followed by Coomassie blue staining of the gel. An aliquot of the purified MP library (∼100 μg; ∼16 μL) was treated with 6 M urea at room temperature for 30 min, followed by reduction with 10 mM DTT for 1 hour. Alkylation was performed with 20 mM IAA in the dark at room temperature for 30 min, followed by re-addition of 10 mM DTT for 30 min. The urea concentration was diluted to 1 M with 50 mM ammonium bicarbonate, pH 8.0, and trypsin was added at an enzyme/protein ratio of 1:100 for 24 h at 25°C on a shaker. Subsequently, the digested peptides were acidified to pH 3 with the addition of ∼20μl 10% formic acid and desalted using hand-packed Stage-Tips C18. The eluted peptides were dried by vacuum centrifugation.

### LC and MS/MS analyses

NanoLC connected to an Orbitrap Exploris mass spectrometer (Thermo Fisher Scientific) was used for the analysis of all samples. The peptide separation was carried out using a Proxeon EASY nLC 1200 System (Thermo Fisher Scientific) fitted with a custom-made C18 column (15 cm x 150 μm ID) packed with HxSil C18 3 μm Resin 100 Å (Hamilton). A gradient of water/acetonitrile/0.1% formic acid was employed for chromatography. The samples were injected onto the column and run for 180 minutes at a flow rate of 0.60 μl/min. The peptide separation began with 1% acetonitrile, increasing to 3% in the first 4 minutes, followed by a linear gradient from 3% to 23% acetonitrile over 86 minutes, then another increase from 24% to 80% acetonitrile over 35 minutes, and finally a 35-minute wash at 80% acetonitrile, and then decreasing to 1% acetonitrile for 10 min and kept 1% acetonitrile for another 10 min. The eluted peptides were ionized using positive nanoelectrospray ionization (NSI) and directly introduced into the mass spectrometer with an ion source temperature set at 250°C and an ion spray voltage of 2.1 kV. All of the full-scan MS spectra (m/z 375-1500) were captured at an Orbitrap in the resolution 15000 with the defined m/z range 145-1450 m/z. The Orbitrap Exploris mass spectrometer was operated using Thermo XCalibur software.

### Data Analysis in DIA-NN

Data were processed using DIA-NN version 1.9.1 in library-free mode with contaminants enabled. FASTA digest–based library generation was performed with deep learning–based prediction of spectra, retention times (RTs), and ion mobilities (IMs) using the *Mus musculus* reference proteome (UniProt proteome ID: UP000000589). Trypsin/P was specified as the protease, allowing a maximum of one missed cleavage. No variable modifications were permitted, while N-terminal methionine excision and cysteine carbamidomethylation were enabled. The peptide length range was set to 7–30 amino acids, with a precursor charge range of 1–4. The precursor m/z range was 300–1800, and the fragment ion m/z range was 200–1800.

Quantification matrices were enabled, with precursor-level false discovery rate (FDR) controlled at 1% and the log level set to 1. Heuristic protein inference was enabled with shared spectra excluded, and protein inference was performed at the gene level. The neural network classifier was run in single-pass mode. Quantification was performed using the QuantUMS (high-precision) strategy, with RT-dependent cross-run normalization enabled. Library generation included peptide identifications, RT, and IM profiling. Speed and RAM usage were set to optimize identification and quantification performance. All MS datasets were processed together within a single DIA-NN run to ensure consistent identification, quantification, and cross-run normalization across all samples.

### Protein annotation

The protein list obtained from DIA-NN was subjected to a gene ontology (GO)-term analysis using the UniProtKB database to identify proteins with the GO-term “membrane”. Within this group, proteins containing at least one α-helical transmembrane segment were labeled as IMPs. While those without any transmembrane segment were annotated as MAPs. The Phobius web server was utilized (http://phobius.sbc.su.se/) to predict the number of transmembrane segments (44). The subcellular localization of the IMPs was further classified using the GO-term “Subcellular location [CC].” These were divided into pIMPs if they contain the GO-term ‘plasma membrane’. Alternatively, they were labeled as oIMPs when containing keywords such as ER, Golgi membrane, vesicle membrane (including endosome, exosome, peroxisome, lysosome, and vesicle), mitochondrial and nucleus membranes).

### Statistical analysis

The report.PG_matrix.tsv output generated by DIA-NN was imported into Perseus (v1.6.15.0) for downstream statistical analysis. LFQ intensity values were log₂-transformed before analysis. Differential protein capture between disease state or VU0486846 treatment conditions was assessed using a two-sample Student’s *t*-test with a within-group variance parameter (*s*₀) set to 0.1. Proteins were retained for statistical testing only if at least three valid intensity values were present in either the treatment or control group. Missing values were imputed from a normal distribution downshifted by 1.8 standard deviations from the global mean with a width of 0.3 times the standard deviation. Proteins were considered significantly differentially enriched if they met thresholds of −log₁₀(*p*-value) > 1.3 and an absolute log₂ fold change greater than 0.5. All subsequent data analysis and visualization were performed in RStudio (v4.5.1).

## Results

### Membrane Proteome Sample Preparation from Mouse Cortex

The membrane proteomics workflow employed here has been previously published and validated for the enrichment and quantitative analysis of integral membrane proteins IMPs (12). Briefly, crude membranes were isolated from the cerebral cortex of female B6C3F1 mice by French press homogenization in a hypotonic buffer containing protease inhibitors and DNase, followed by differential centrifugation and ultracentrifugation (110,000 x g). Membranes were processed in biological triplicate across four mouse groups (WT vehicle, WT VU0486846, APP vehicle, and APP VU0486846). IMPs were solubilized with 1% DDM and reconstituted using the Peptidisc scaffold before purification by Ni–NTA affinity chromatography. The workflow consistency across biological replicates was verified by SDS–PAGE, which revealed highly similar protein profiles within each experimental group (**Figure 1A**). Equal quantities of each sample were then subjected to LC–MS/MS analysis.

**Figure 1.**
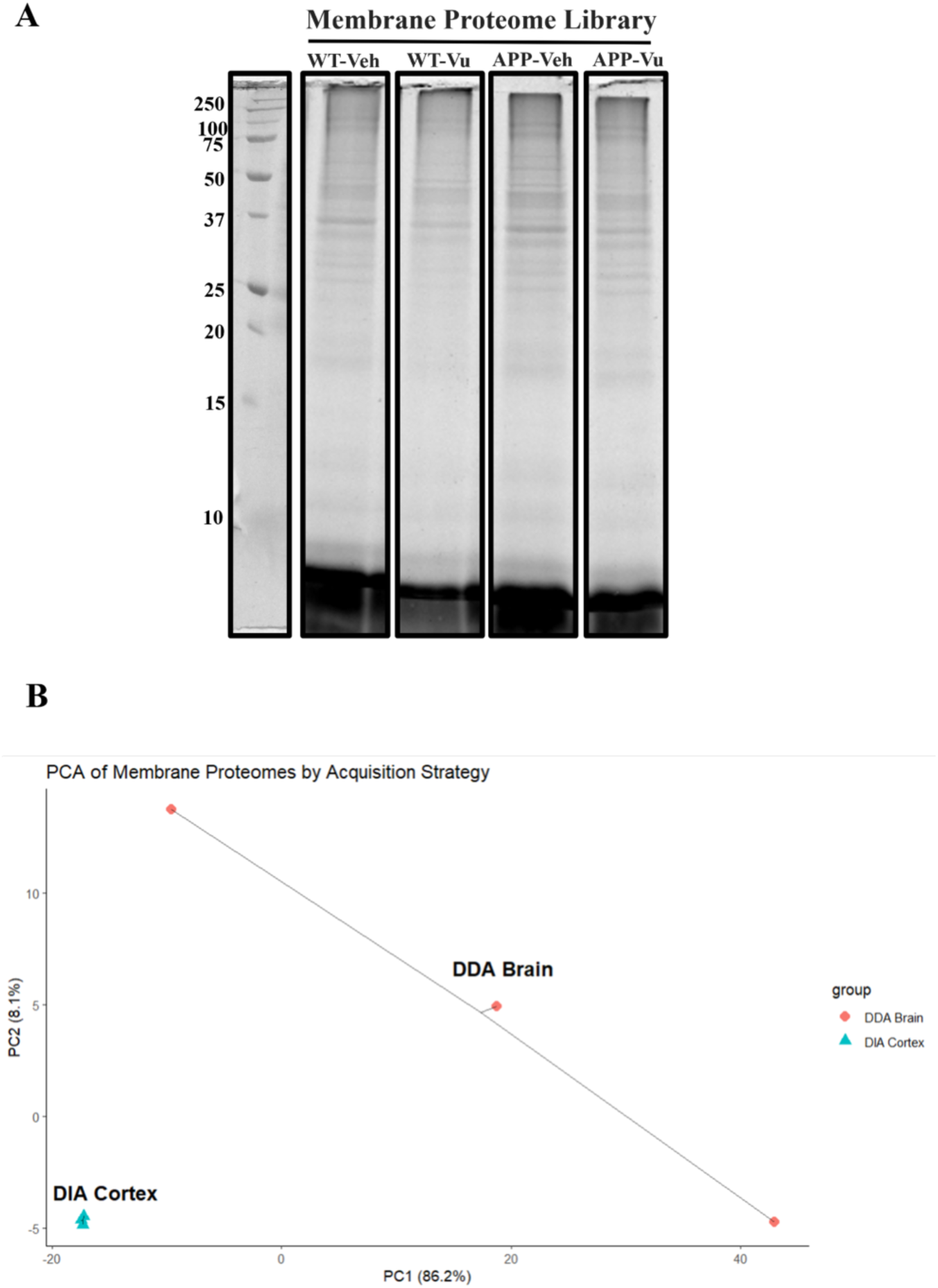
Peptidisc-based membrane proteome libraries and acquisition-dependent differences between DDA mouse brain and DIA mouse cortex datasets. (A) SDS–PAGE analysis of peptidisc-reconstituted mouse cortex membrane proteome libraries analyzed in this study. WT-Veh: Wild-type mouse treated with vehicle; WT-Vu: Wild-type mouse treated with VU0486846; APP-Veh: APP mouse treated with vehicle; APP-Vu: APP mouse treated with VU0486846. Molecular weight marker on the left part of the gel. (B) Principal component analysis (PCA) comparing DIA-acquired wild-type mouse cortex datasets with DDA-acquired mouse brain datasets. The plot was generated with the ggrepel packages in r.

### Comparison of DDA and DIA for Membrane Proteome Profiling

Our peptidisc-enabled membrane proteomics platform has been developed and validated using data-dependent acquisition (DDA) mass spectrometry in multiple prior studies(12,20–23). However, recent advances in acquisition strategies and data analysis have driven the widespread adoption of data-independent acquisition (DIA), which offers improved quantitative reproducibility, depth of coverage, and consistency across large and complex proteomic datasets (24–26). Here, to benchmark this DIA-based acquisition to membrane proteomic profiling, we compared the DIA dataset generated from WT mouse cortex in the present study with our previously published DDA-based mouse brain membrane proteome dataset (20). This comparison enabled us to assess how the acquisition strategy influences the detection and quantification of IMPs, independent of disease state.

We first evaluated global variance structure and inter-replicate consistency using principal component analysis (PCA). PCA revealed that the DDA dataset exhibited the greatest inter-replicate variability, whereas the DIA dataset showed tight clustering across replicates (**Figure 1B**), indicating improved quantitative reproducibility with DIA. Analysis of protein counts further showed that DDA identified the fewest IMPs overall (465 ± 68) compared with DIA (840 ± 7; Table 1). In contrast, the DDA dataset exhibited a higher relative enrichment of IMPs (48.2% ± 7.2%) than the DIA dataset (38.4% ± 0.3%), suggesting that DDA preferentially captures highly abundant IMPs. These observations indicate that differences in replicate variability are closely linked to differences in proteome depth and abundance bias. In agreement with this interpretation, analysis of protein intensity distributions showed that DDA preferentially detected integral membrane proteins among the most abundant species, with 45.5% of the top 200 proteins classified as IMPs in the DDA dataset compared with 38.5% in the DIA dataset (**Figure 2A**).

**Figure 2.**
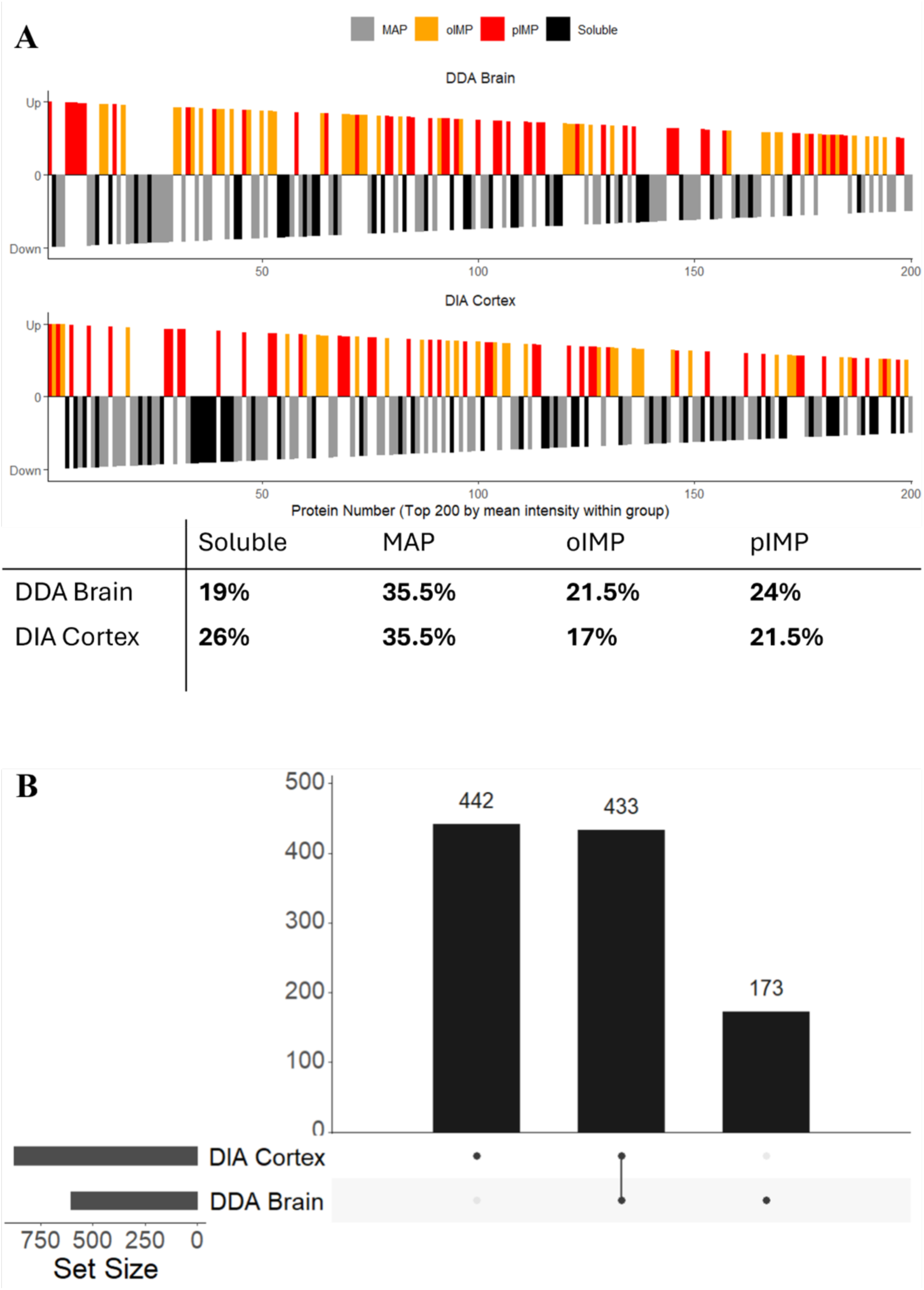
Comparative enrichment and overlap of membrane proteins in DDA and DIA datasets. (A) Distribution of the top 200 most abundant proteins in the DIA mouse cortex and DDA mouse brain datasets, with an accompanying table summarizing relative enrichment by subcellular localization. MAP: membrane-located proteins. oIMPs: organelle-located integral membrane proteins. pIMPs: plasma membrane-located integral membrane proteins. Soluble: soluble proteins. Localization percentages were calculated from the top 200 proteins ranked by replicate-averaged intensities within each dataset. As these values represent the composition of a defined protein set rather than a distribution of replicate measurements, standard deviations are not reported. Plotted using the ggplot2 package in r. (B) UpSet plot illustrating the overlap of IMPs identified in the DIA cortex and DDA brain datasets. The analysis was performed in R using the Complex Upset package.

**Table 1.** Comparison of protein identifications between wild-type DDA mouse brain and DIA mouse datasets.

|  | TP | SP | MAP | tIMP | oIMP | pIMP | %<br>tIMPs | %<br>pIMPs |
| --- | --- | --- | --- | --- | --- | --- | --- | --- |
| DDA<br>mBrain | 963<br>+/- 141 | 498<br>+/- 73 | 308<br>+/- 43 | 465<br>+/- 68 | 240<br>+/- 37 | 225<br>+/- 31 | 48.2%<br>+/- 7.2% | 23.4%<br>+/- 3.2% |
| DIA<br>mCortex | 2190<br>+/- 27 | 1349<br>+/- 21 | 711<br>+/- 6 | 840<br>+/- 7 | 425<br>+/- 4 | 416<br>+/- 4 | 38.4%<br>+/- 0.3% | 19%<br>+/- 0.2% |

To determine how these differences in reproducibility and abundance bias translate into protein-level overlap and method-specific detection, we next performed UpSet plot analysis. This analysis revealed both substantial overlap and clear difference between the datasets, with 433 shared IMPs, 442 unique to the DIA cortex dataset, and 173 unique to the DDA brain dataset (**Figure 2B**). Furthermore, in our previous mouse multi-organ DDA study, we identified 121 pIMPs enriched in the brain relative to four other organs. Of these, 118 were also detected in the WT DIA cortex dataset in the present study, demonstrating strong concordance between acquisition strategies for highly abundant, brain-enriched IMPs. This finding indicates strong concordance between acquisition strategies for highly expressed IMPs, despite differences in tissue composition, while highlighting the expanded detection of lower-abundance IMPs achieved with DIA.

Altogether, both methods robustly quantify abundant IMPs, but DIA provides greater proteome depth and reproducibility, whereas DDA preferentially detects highly expressed IMPs and underrepresents lower-abundance proteins. Consequently, DIA emerges as better suited for unbiased, discovery-driven membrane proteome profiling in a disease context when targets are not known *a priori*.

### Assessment of Global Structure in DIA Membrane Proteome Datasets

Because this study examines two experimental variables, the APP genotype and M1 mAChR modulation, we structured our analyses to separately assess genotype-dependent effects and VU0486846-driven effects. However, as an initial quality-control step, we evaluated dataset quality and reproducibility by comparing total protein identifications across all conditions. Herein, a consistent number of IMPs (802-841, representing 38–39% of total identifications, **Table 2**), with no differences in protein IDs attributable to genotype or VU0486846 treatment.

**Table 2.** Protein count identifications of DIA-MS datasets analyzed in this study.

|  | TP | SP | MAP | tIMP | oIMP | pIMP | %<br>tIMPs | %<br>pIMPs |
| --- | --- | --- | --- | --- | --- | --- | --- | --- |
| WT Vehicle | 2190<br>+/- 27 | 1349<br>+/- 21 | 711<br>+/- 6 | 841<br>+/- 7 | 425<br>+/- 4 | 416<br>+/- 4 | 38.4%<br>+/- 0.3% | 19%<br>+/- 0.18% |
| WT VU0486846 | 2145<br>+/- 38 | 1323<br>+/- 24 | 703<br>+/- 13 | 823<br>+/- 14 | 417<br>+/- 8 | 406<br>+/- 6 | 38.3%<br>+/- 0.65% | 18.9%<br>+/- 0.27% |
| APP Vehicle | 2061<br>+/- 36 | 1260<br>+/- 18 | 681<br>+/- 5 | 802<br>+/- 18 | 408<br>+/- 11 | 393<br>+/- 8 | 38.9%<br>+/- 0.87% | 19.1%<br>+/- 0.39% |
| APP VU0486846 | 2140<br>+/- 55 | 1308<br>+/- 41 | 700<br>+/- 11 | 832<br>+/- 14 | 421<br>+/- 6 | 411<br>+/- 9 | 38.9%<br>+/- 0.65% | 19.2%<br>+/- 0.42% |

We next performed PCA to examine the overall dataset structure. WT and APP vehicle-treated samples showed tight clustering, whereas VU0486846-treated samples, particularly when combined with genotype effects, exhibited increased variance, including a single outlier (**Figure 3A**). Although all datasets exhibit comparable protein identification counts and overall proteome coverage, indicating uniform data quality, PCA reveals increased dispersion among ligand-treated samples. This suggests that the introduction of VU0486846 increases biological variability between replicates without compromising dataset integrity.

**Figure 3.**
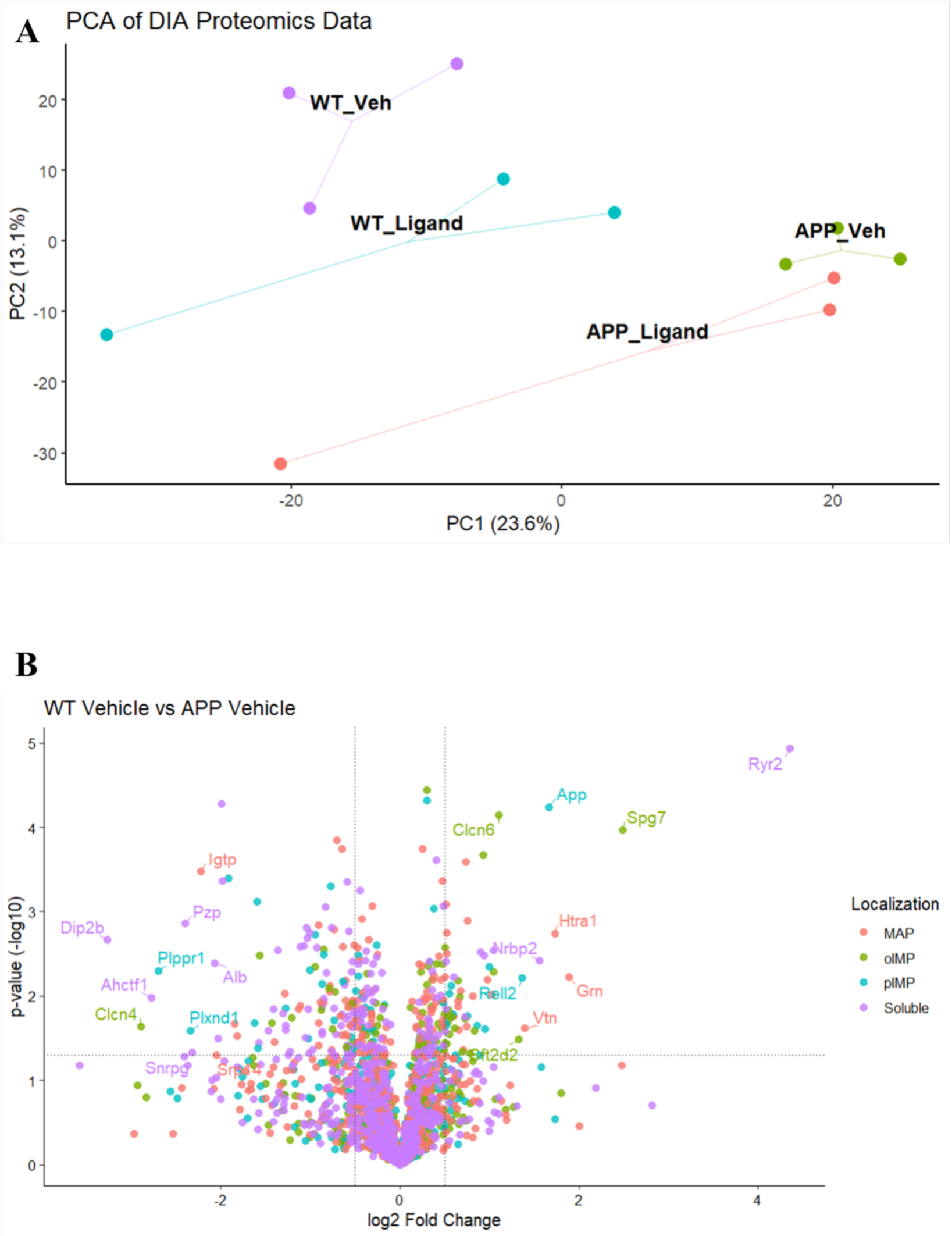
Global structure of DIA datasets and differential membrane protein capture in WT and APP Mice. (A) Principal component analysis of DIA-acquired membrane proteome datasets analyzed in this study. WT_Veh: Wild-type mouse treated with vehicle; WT_Vu: Wild-type mouse treated with VU0486846; APP_Ve: APP mouse treated with vehicle; APP_Vu: APP mouse treated with VU0486846. The plot was generated with the factoextra and ggrepel packages in r. (B) Volcano plot comparing 3 vehicle-treated wild-type and 3 vehicle-treated APP mice, with proteins colored by subcellular localization. MAP: Membrane-located proteins. oIMPs: Organelle-located integral membrane proteins. pIMPs: Plasma membrane-located integral membrane proteins. Soluble: Soluble proteins. A two-tailed *t* test was performed in Perseus with *s0* = 0.1, and significance thresholds were set at log2 FC ≥ ±0.5 and –log10(*p*-value) ≥ 1.3. Transmembrane domains were predicted using Phobius to classify proteins as pIMPs (*blue*) or oIMPs (*green*). Only proteins with valid intensity values in at least three out of six replicates were included, with missing values imputed using a width of 0.3 and a downshift of 1.8. Plots generated using the ggplot2 package in R.

We next quantified the number of differentially captured IMPs across all pairwise comparisons (**Table 3**). Comparison of WT versus APP mice revealed 64 differentially expressed IMPs (7.8% of all IMPs), whereas only 8 were detected between vehicle- and VU0486846-treated in WT mice (0.98% of all IMPs) and 15 between vehicle- and VU0486846-treated in APP mice (3.4% of all IMPs). Together, these results indicate that APP genotype exerts a substantially larger effect on the global membrane proteome than M1 modulation with VU0486846.

**Table 3.** Number of differentially captured proteins as determined by Volcano plot comparisons performed in this study.

| Comparison | Number of<br>IMPs in<br>comparison | Number IMPs<br>enriched in<br>condition 1 | Number IMPs<br>enriched in<br>condition 2 | % of IMPs<br>Differentially<br>captured |
| --- | --- | --- | --- | --- |
| WT vs. App | 813 | 32 | 32 | 7.8% |
| WT vs. WT<br>ligand | 818 | 5 | 3 | 0.98% |
| APP vs APP<br>ligand | 802 | 6 | 22 | 3.4% |

### Global APP-Driven Reorganization of the Cortical Membrane Proteome

Comparison of WT and APP vehicle-treated mice revealed substantial dysregulation of the cortical membrane proteome associated with AD progression. Volcano plot analysis identified 64 significantly altered IMPs, with 32 enriched in WT and 32 enriched in APP mice, out of 813 quantified IMPs, corresponding to 7.8% of the membrane proteome (**Figure 3B**). This extent of dysregulation is consistent with the widespread membrane-associated remodeling reported in AD (6,9,27).

We first applied Gene Ontology (GO) enrichment analysis to assess global pathway-level perturbations associated with APP pathology. In the WT versus APP comparison, IMPs were enriched for pathways related to inflammatory response, ion transport, membrane organization, and nervous system development (**Supplementary Figure 1A**), consistent with broad remodeling of membrane-associated homeostatic and immune processes during AD progression (28,29). In contrast, APP-biased IMPs showed significant enrichment for synapse organization, cell–cell adhesion, and cell junction pathways (**Supplementary Figure 1B**), reflecting disruption of synaptic and structural membrane organization in the APP cortex (28,30). As expected, APP itself was among the most significantly enriched proteins in APP mice, exhibiting the strongest statistical signal among all differentially detected IMPs (-log10 *p* = 4.24, log2 fold change = 1.67 to APP; *i.e* 3.2-fold increase). This result is consistent with transgenic APP overexpression driving pathology in this model and serves as an internal validation of the membrane-focused Peptidisc workflow.

### APP-Enriched Membrane Proteins Highlight Dysregulated Calcium Signaling, Endolysosomal Trafficking, and Synaptic Membrane Homeostasis

We next examined the functional identity of IMPs that are enriched in the APP mice. Two endolysosomal IMPs, PLD3 (−log10 *p* = 3.67; log2 fold change = 0.926; ∼1.9-fold) and ITM2C (−log10 *p* = 1.78; log2 fold change = 0.587; ∼1.5-fold) were significantly enriched. PLD3 is an AD risk factor whose reported abundance changes in the human AD brain have been variable, suggesting context-dependent or compensatory regulation (27,28). In contrast, ITM2C, a close homolog of the familial dementia–associated protein ITM2B (BRI2), has been reported to have elevated abundance in AD, consistent with its role in amyloid precursor protein trafficking (33,34). The enrichment of both PLD3 and ITM2C observed here supports dysregulation of the endolysosomal pathway as a prominent feature of APP-driven pathology (35).

CNTNAP2, which encodes the neurexin-family cell adhesion molecule CASPR2, was also enriched in APP mice (−log10 *p* = 1.74; log2 fold change = 0.724; ∼1.65-fold). CASPR2 localizes to axonal and synaptic membranes and plays established roles in neuronal connectivity and synaptic organization (36). CNTNAP2 has been identified as a potential AD susceptibility locus, and transcriptomic studies reporting reduced expression in affected brain regions, suggesting complex and context-dependent regulation across disease stages and experimental systems(37,38). Its enrichment in the APP cortex further highlights the disruption of membrane-associated adhesion and connectivity pathways during disease progression.

Notably, a soluble partial fragment of ryanodine receptor 2 (RyR2) emerged as the most significantly differentially captured protein, exhibiting strong enrichment in APP mice (−log10 *p* = 4.93; log2 fold change = 4.35; ∼20-fold). Aberrant RyR2-mediated calcium release has been linked to heightened neuronal excitability and accelerated disease progression in AD models, and genetic modulation of RyR2 gating has been shown to ameliorate cognitive deficits in familial AD mice (39,40). The pronounced enrichment of RyR2-derived species observed here is therefore consistent with dysregulated calcium signaling as a key feature of APP-driven pathology (41).

In addition, glial fibrillary acidic protein (GFAP), a soluble astrocytic marker, was significantly enriched in APP mice (−log10 *p* = 1.59; log2 fold change = 0.656). Although GFAP is not an IMP, its association with membrane-proximal cytoskeletal structures likely explains its recovery in our membrane-focused workflow. This finding is consistent with our prior report of increased GFAP expression in hippocampal lysates from APP/PS1 mice (15) and reflects the ability of the peptidisc-based profiling to capture membrane-proximal astrocytic components accompanying disease progression.

Collectively, these results highlight that AD pathology in APP mice is associated with coordinated remodeling of the cortical membrane proteome. Proteins enriched in APP mice include several well-established AD–associated membrane proteins, such as Ryr2, PLD3, ITM2C, and CNTNAP2, consistent with prior genetic and proteomic studies implicating calcium signaling, endolysosomal trafficking, and synaptic membrane dysfunction in disease pathogenesis.

### Disruption of Neuronal Maintenance Pathways in APP Mouse Cortex

Among proteins depleted in APP mice, several clustered into pathways governing axon guidance and neuronal connectivity, including EPHA5, ROBO2, PLXND1, PLXNB2, CDH10, and KIT. These proteins are components of Eph receptor, Slit–Robo, and plexin–semaphorin signaling pathways that regulate axon pathfinding, synaptic organization, and circuit maintenance in the mature brain. All three of these pathways have been previously implicated in AD pathology and corroborate our findings with AD-implicated proteins (42–44).

In addition, multiple proteins involved in synaptic organization and plasticity were reduced in the APP cortex, including NFASC, ELFN1, CADM1, ALCAM, GRM2, P2RY6, and SYT12. These proteins participate in synaptic adhesion, neurotransmitter signaling, and activity-dependent synaptic modulation, processes that are critical for synaptic stability and cognitive function. Their coordinated depletion in APP mice is consistent with extensive evidence that synaptic adhesion molecules and glutamatergic signaling pathways are disrupted in AD and contribute to synaptic dysfunction and cognitive decline (45–47).

Proteins involved in cytoskeletal organization were likewise depleted in APP mice, including SPAST, PTPN5, and ARHGAP26. The latter is a Rho GTPase-activating protein that regulates RhoA-dependent actin dynamics and has been genetically linked to AD risk, with prior studies reporting associations between ARHGAP26 polymorphisms and AD susceptibility (48). Loss or dysregulation of cytoskeletal regulators such as spastin (SPAST) and STEP/PTPN5 has been associated with neurite degeneration, synaptic weakening, and cytoskeletal instability in AD and related tauopathies(49,50). The coordinated depletion of these proteins in APP mice is therefore consistent with the broader literature showing microtubule and actin cytoskeletal disruption during AD progression.

Finally, FAM234B, a poorly characterized neuron-enriched IMP, was depleted in APP mice. Although its biological function remains largely undefined, FAM234B is a close paralog of FAM234A, which has been reported to be dysregulated in neurodegenerative contexts, including AD–associated epigenomic studies (51,52). Its selective reduction in the APP cortex highlights the ability of this membrane-centric workflow to capture changes in understudied membrane proteins and suggests FAM234B as a potential candidate for further investigation in AD.

Together, IMPs depleted in APP mice predominantly map to pathways governing axon guidance, synaptic organization and plasticity, and cytoskeletal regulation, highlighting the progressive loss of membrane-associated programs required for neuronal maintenance and connectivity during AD progression.

### Effect of VU0486846 on the WT Mice Global Membrane Proteome

In the comparison between WT vehicle– and VU0486846-treated WT mice, only a small number of IMPs exhibited significant differential expression. Five IMPs were enriched in vehicle-treated WT samples (DGKE, PTPRZ1, YIF1B, DNER, and LPAR1), while three were enriched following VU0486846 treatment (RELL2, CDH16, and SPG7), out of 818 quantified IMPs (**Figure 4A**). Thus, fewer than 1% of the WT membrane proteome was altered by VU0486846 treatment, indicating minimal global membrane remodeling, particularly when contrasted with the extensive membrane dysregulation observed in APP mice.

**Figure 4.**
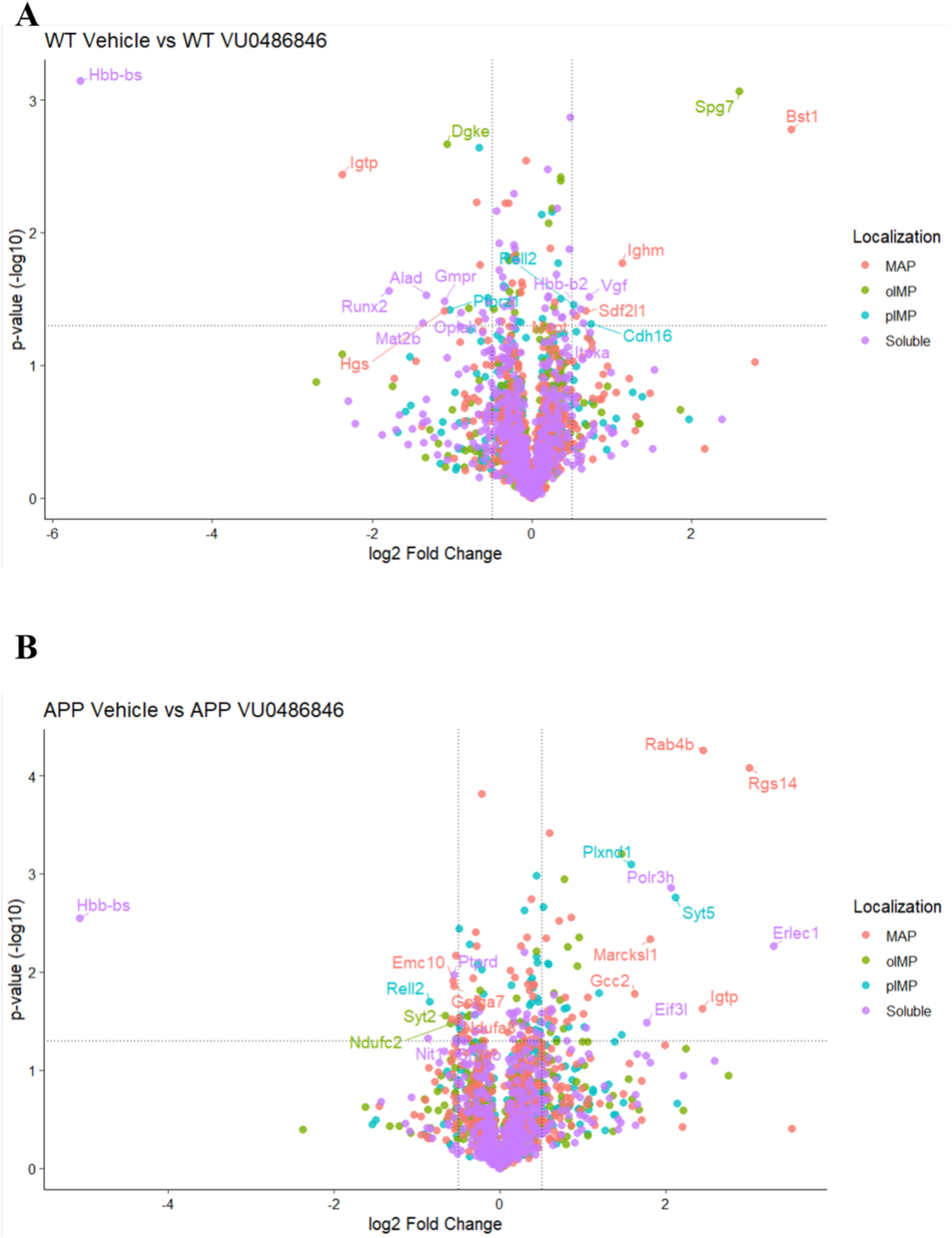
Differential membrane protein capture following VU0486846 treatment. (A) Volcano plot comparing 3 vehicle-treated wild-type mice with 3 wild-type mice treated with VU0486846. MAP: membrane-located proteins. oIMPs: organelle-located integral membrane proteins. pIMPs: plasma membrane-located integral membrane proteins. Soluble: soluble proteins. A two-tailed *t* test was performed in Perseus with *s0* = 0.1, and significance thresholds were set at log2 FC ≥ ±0.5 and –log10(*p*-value) ≥ 1.3. Transmembrane domains were predicted using Phobius to classify proteins as pIMPs (*blue*) or oIMPs (*green*). Only proteins with valid intensity values in at least three out of six replicates were included, with missing values imputed using a width of 0.3 and a downshift of 1.8. Plots generated using the ggplot2 package in R. (B) Volcano plot comparing 3 vehicle-treated APP mice with 3 APP mice treated with VU0486846. Statistical analysis and plotting same as A.

Among the WT-enriched proteins, PTPRZ1 and LPAR1 are Gq-coupled signaling proteins involved in neuronal plasticity and excitability (53,54). Because activation of the M1 mAChR engages canonical Gq/11-dependent phospholipase C signaling(55), the reduced abundance of these proteins following VU0486846 treatment may reflect engagement of M1-driven Gq signaling, thereby reducing reliance on auxiliary Gq-coupled pathways.

The remaining differentially captured IMPs lack established links to M1 mAChR signaling and likely reflect minor, context-dependent differences in IMP capture rather than coordinated pathway-level engagement. Collectively, these findings indicate that pharmacological M1 mAChR activation in WT mice produces minimal changes in the global membrane proteome.

### Effect of VU0486846 on APP Mice Global Membrane Proteome

Comparison of vehicle- and VU0486846-treated APP mice revealed modest but informative changes in the membrane proteome. Of 818 quantified IMPs, 6 and 22 were enriched in vehicle- and VU0486846-treated mice, respectively (3.4% overall IMPs differentially enriched; **Figure 4B**), indicating that M1 mAChR activation induces selective membrane remodeling rather than broad, nonspecific changes. Go term enrichment analysis showed in VU0486846-treated APP mice, enriched GO terms included regulation of biological quality, transport, synapse organization, neuron recognition, and cell junction organization (**Supplementary Figure 1C**). These pathway-level changes are consistent with selective modulation of synaptic, trafficking, and cell-cell interaction processes following M1 mAChR activation (52–55).

Among the IMPs enriched in VU0486846-treated APP mice, several mapped to pathways governing neuronal trafficking, axon guidance, and synaptic organization, including SORCS2, EPHA5, PLXND1, PLXNB2, CADM1, and PCDH10. The protein SORCS2 regulates intracellular trafficking, EPHA5 and the plexin receptors PLXND1 and PLXNB2 control axon guidance and synaptic patterning, while CADM1 and PCDH10 support synapse formation and stability, processes that are disrupted during AD progression (60–65). The increased abundance of these proteins following VU0486846 treatment is therefore consistent with modulation of disease-associated synaptic trafficking and organization.

Consistent with engagement of canonical M1 mAChR signaling, GPRIN1 and RASAL1, two proteins directly linked to Gq-dependent pathways, were also enriched following VU0486846 treatment. GPRIN1 is a membrane-associated protein that interacts with Gαq/11 and functions as a downstream effector of Gq-coupled GPCR signaling, with established roles in neurite outgrowth and synaptic plasticity (66). RASAL1 is a Ras GTPase-activating protein that negatively regulates Ras–ERK signaling, a pathway downstream of Gq activation and known to be dysregulated in AD (67). Their enrichment in the VU0486846-treated APP mice is consistent with compensatory upregulation of downstream signaling components following M1 mAChR activation.

In contrast, proteins enriched in vehicle-treated APP mice (RELL2, SYT2, NDUFC2, KCNC3, NKG7, and STX1B) are not established components of M1 mAChR signaling but instead map to synaptic (SYT2, STX1B), excitability (KCNC3), mitochondrial (NDUFC2), and immune (NKG7) pathways implicated in AD-associated network dysfunction (68–74). Their reduction following VU0486846 treatment is therefore consistent with downstream, disease-state–dependent remodeling rather than direct receptor-proximal effects.

Collectively, these findings indicate that M1 mAChR activation in APP mice induces targeted remodeling of IMPs involved in neuronal trafficking, synaptic organization, and intracellular signaling, while avoiding broad perturbations of the membrane proteome. This targeted pattern of change is consistent with improved neuronal function rather than nonspecific off-target effects, supporting the therapeutic relevance of M1 mAChR positive allosteric modulation in AD models.

## Discussion

Because AD pathology is driven by dysregulation of membrane-associated signaling, transport, and adhesion processes(75), proteomic strategies that enrich and preserve native IMP assemblies are essential. In contrast, conventional bulk proteomic approaches rely on denaturing extraction conditions that limit resolution of membrane-embedded receptors, ion channels, and trafficking systems (76), resulting in systematic underrepresentation of disease-relevant IMPs.

Accordingly, prior AD proteomic network analyses of brain and cerebrospinal fluid have been dominated by soluble metabolic enzymes, complement components, and glial markers, with relatively few IMPs identified (8), and a recent integrative analysis of human and mouse AD datasets reported only 9 IMPs among 37 human-specific differentially expressed proteins (77).

To address this limitation, we applied peptidisc-enabled, DIA-based membrane proteomics to define how the cortical membrane proteome is remodeled during AD progression and in response to M1 mAChR activation. Benchmarking our WT DIA mouse cortex dataset against our previously published DDA mouse brain dataset showed that DIA provides improved quantitative reproducibility and greater coverage of IMPs, supporting its use for disease-state membrane proteome analysis. This finding is consistent with the increasing adoption of DIA-based workflows for complex biological systems.

Using this approach, we recovered numerous APP-enriched IMPs with known AD-associations such as RyR2, PLD3, ITM2C, and CNTNAP2. Notably, a previous membrane enrichment–based MS study of post-mortem AD brain tissue identified only 13 differentially enriched proteins, none of which were IMPs, likely reflecting solubilization of crude membrane fractions using urea without detergent extraction or membrane-mimetic stabilization (78). In contrast, the peptidisc-based workflow employed here enabled detection of 64 differentially captured IMPs between WT and APP mouse cortex, underscoring the improved ability of membrane-mimetic proteomics to resolve disease-associated membrane protein dysregulation.

Within this expanded IMP landscape, RyR2 emerged as the most strongly differentially captured protein, albeit in an unexpected form. The dominant signal corresponded not to full-length RyR2, but to a soluble fragment (protein ID A0A1Y7VK09). Examination of the peptide-level report.pr_matrix revealed that when all RyR2-derived peptides were grouped into a single canonical protein group, no significant differential enrichment was observed, indicating that the observed change was restricted to the fragment-associated peptide rather than reflecting altered total RyR2 abundance. Altered RyR2 gating and enhanced ER-to-cytosol Ca²⁺ leak have been implicated in AD pathophysiology and neuronal hyperactivity (79,80). Thus, it is plausible that AD-associated RyR2 dysregulation alters channel conformation or post-translational modification state, increasing the accessibility or stability of specific cytosolic RyR2 regions and resulting in preferential detection of fragment-specific peptides without a corresponding increase in overall channel abundance.

At the systems level, APP pathology was associated with extensive remodeling of the cortical membrane proteome, with 7.8% of detected IMPs differentially enriched between WT and APP mouse cortex. In contrast, proteins enriched in WT mice predominantly reflected homeostatic and neuronal maintenance pathways, including axon guidance, cytoskeletal organization, and synaptic integrity. Modulation of M1 mAChR signaling induced substantially fewer global membrane changes (0.98% in WT mice and 3.4% in APP mice) and selectively increased the abundance of proteins involved in synaptic organization, trafficking, and neuronal structure, consistent with engagement of membrane pathways linked to neuronal homeostasis.

Taken together, the contrast between extensive APP-driven membrane remodeling and the more selective effects of M1 mAChR modulation highlights a broad dynamic range of membrane proteome responses, demonstrating that peptidisc-enabled membrane proteome profiling has sufficient sensitivity to distinguish large-scale pathological perturbations from subtler, context-dependent pharmacological effects. This level of resolution enables longitudinal and comparative analyses aimed at tracking temporal changes in membrane proteome organization and distinguishing primary disease drivers from secondary or compensatory responses. This capability is particularly relevant given the heterogeneity of AD mouse models, which vary widely in disease onset, progression, cellular vulnerability, and pathway engagement (81). The sensitivity and reproducibility observed here suggest that peptidisc-based membrane proteomics can be broadly applied across models to define model-specific membrane signatures and assess how distinct genetic drivers reshape IMP networks, thereby informing model selection for targeted biological questions. In this study, whole-cortex profiling was prioritized to ensure robust and reproducible membrane protein recovery, as current peptidisc workflows require substantial input material, however continued methodological advances are expected to enable extension of this approach to region- and subregion-specific analyses.

In conclusion, this study establishes peptidisc-enabled, DIA-based membrane proteomics as a robust and sensitive platform for defining disease-associated remodeling of the native membrane proteome. By enriching and preserving IMPs from brain cortex tissue, we captured extensive APP-driven membrane dysregulation alongside more selective, context-dependent changes induced by M1 mAChR modulation. Together, these findings position membrane-centric proteomics as a powerful strategy for elucidating IMP-level mechanisms of neurodegeneration and for identifying membrane-associated biomarkers and therapeutic targets in AD.

## Supplemental Data

This article contains supplemental data. Supplemental figure (word document containing the supplemental figure). Supplementary file (excel sheet containing the raw data used to make figures).

## Notes

The authors declare the following competing financial interest(s): FDVH is the scientific founder of Peptidisc Biotech. The mass spectrometry proteomics data have been deposited to the ProteomeXchange Consortium via the PRIDE partner repository with the data set identifier PXD074065. Reviewer access details Log in to the PRIDE website using the following details: Project accession: PXD074065 Token: KOT3SByf2RgJ Alternatively, reviewer can access the dataset by logging in to the PRIDE website using the following account details: Username: Password: UooIskZjmfcV

## Acknowledgements

Work in the Duong lab was supported by the CIHR Project grant PG20R34019. Work in the Babu lab was supported by the Canada Foundation for Innovation and CIHR Foundation grant FDN-154318. A. B. is supported by a UBC 4-year fellowship, an Amplify Doctoral Award from Triangle (Training a new generation of researchers in gastroenterology and liver), and a Mast3 (Mass Spectrometry Team Training and Transition) scholarship. K.S.A-E is a Michael Smith Health Research BC funded Health-Professional Investigator and is also funded by Canadian Institutes of Health Research (CIHR) grant PJT-195977 and a New Investigator grant from the Alzheimer’s Society of Canada. S.S.G.F is funded by CIHR grants PJT-148656 and PJT-165967

---

**Supplemental Figure 1.**
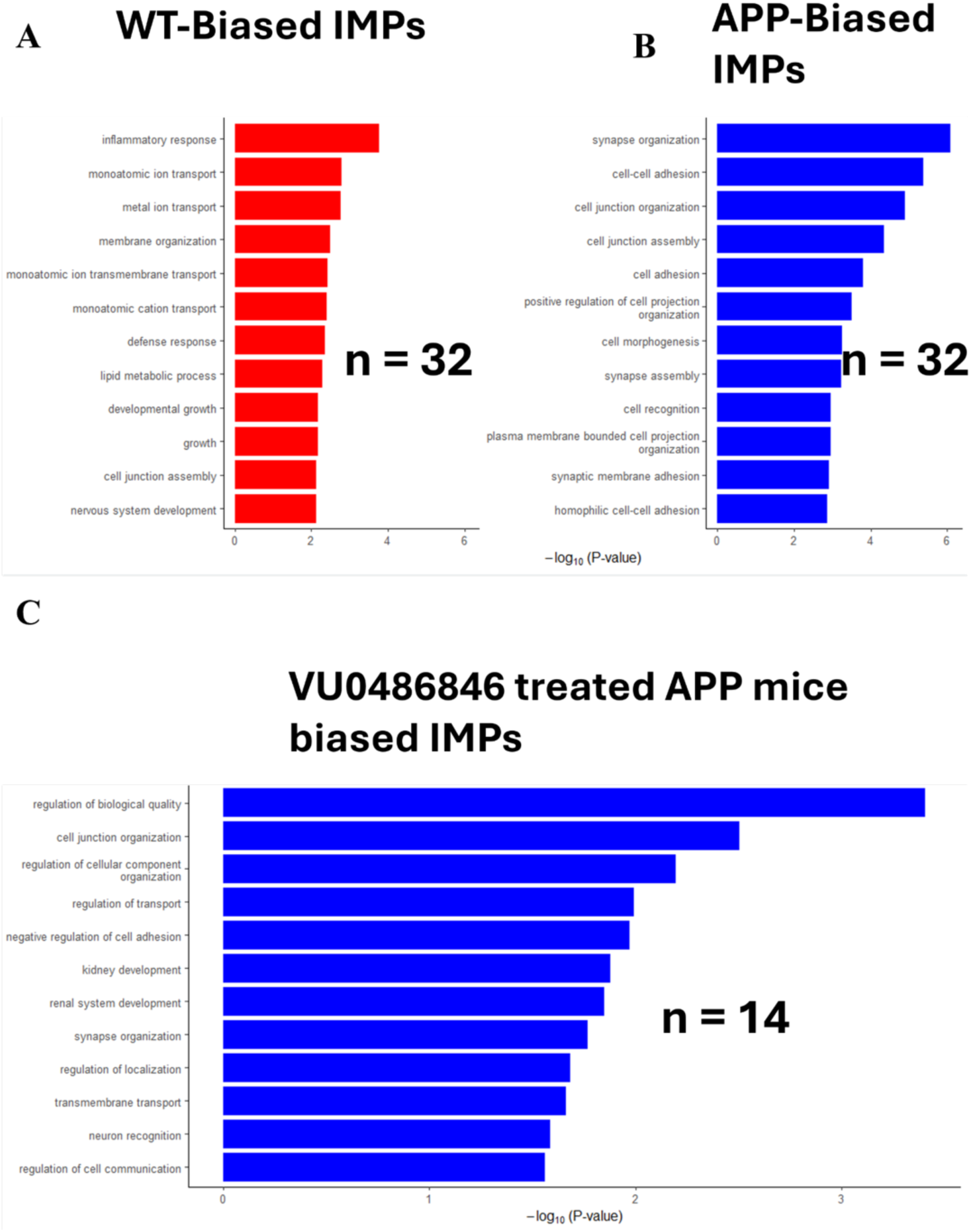
Gene ontology enrichment analysis of differentially captured membrane proteins. (A) Gene Ontology (GO) Biological Process (BP_FAT) enrichment of the top 12 pathways associated with IMPs enriched in WT mice relative to APP mice. Plots were generated in R using ggplot2. (B) GO BP FAT enrichment of IMPs significantly enriched in APP mice relative to wild-type mice. (C) GO BP FAT enrichment of IMPs significantly enriched in VU0486846-treated APP mice relative to vehicle-treated APP mice.

